# Expression Profile of Circular RNAs in Epicardial Adipose Tissue in Heart Failure

**DOI:** 10.1101/764266

**Authors:** Meili Zheng, Lei Zhao, Xinchun Yang

## Abstract

Recent studies have reported circular RNA (circRNA) expression profiles in various tissue types; specifically, a recent work showed a detailed circRNA expression landscape in the heart. However, circRNA expression profile in human epicardial adipose tissue (EAT) remains undefined. RNA-sequencing was carried out to compare circRNA expression patterns in EAT specimens from coronary artery disease (CAD) cases between the heart failure (HF) and non-HF groups. The top highly expressed EAT circRNAs corresponded to genes involved in cell proliferation and inflammatory response, including KIAA0182, RHOBTB3, HIPK3, UBXN7, PCMTD1, N4BP2L2, CFLAR, EPB41L2, FCHO2, FNDC3B and SPECC1. Among the 141 circRNAs substantially different between the HF and non-HF groups (*P*<0.05;fold change>2), hsa_circ_0005565 stood out, and was mostly associated with positive regulation of metabolic processes and insulin resistancein GO and KEGG pathway analyses, respectively. These data indicate EAT circRNAs contribute to the pathogenesis of metabolic disorders causing HF.

## Introduction

Circular RNAs (circRNAs) constitute a new group of non-coding RNAs with a cyclic structure^1^, and contribute to gene regulation. A recent study^2^ carried out deep sequencing of ribosome-free RNAs from 12 human and 25 mousehearts, as well as human embryonic stem cell-derived cardiomyocytesduring differentiation for 28 days, and showed cardiac circRNA expression profile in detail, which provides a valuable reference for the further analysis of circRNAs. Not only have circRNAs been considered intracellular effectors in the pathophysiological alterations of cardiovascular tissues as well as cardiovascular disease markers^3^, they are also involved in multiple cardiovascular ailments^2, 4^.

However, as a key cardio-metabolic factor, epicardial adipose tissue (EAT) is scarcely included in heart tissue specimens in previous studies identifying circRNAs. EAT produces multiple bioactive molecules, including adipokines and cytokines^5, 6^, as well as microparticles carrying proteins, lipids and ribonucleic acids (RNAs)^7^. In addition, EAT is correlated with heart failure(HF), without regard to metabolic status or coronary artery disease (CAD)^8-10^.Meanwhile, EAT releases molecules with vasocrine and paracrine impacts on the myocardium^11, 12^. Therefore, EAT secretome alterations could regulate heart function.

This work aimed to compare circRNA expression patterns in EAT in CAD cases between the HF and non-HF groups. Our findings would supplement circRNA expression patterns in EAT for circRNAdetection in cardiac specimens, also providing plausible HF markers.

## Materials and Methods

### Study Participants

The participants assessed in the present study have been described in our previous report^13^. EAT specimens were obtained from 10 CAD cases undergoing coronary artery bypass grafting in the Department of Heart Center, Beijing Chao-yang Hospital, Capital Medical University. They were assigned to the HF and non-HF groups (n=5/group). HF was defined by brain natriuretic peptide (BNP)>500ng/L and abnormal echocardiograms (left ventricular end diastolic diameter [LVEDD]>50mm and >55mm in females and males, respectively; left ventricular ejection fraction [LVEF]<50%).The non-HF group comprised individuals showing BNP<100 ng/L and normal echocardiograms. This study had approval from the Ethics Committee of Beijing Chao-yang Hospital, Capital Medical University. Each patient provided written informed consent.

### RNA-sequencing Procedure

Figure 1 shows the detailedRNA-sequencing procedure, which was described in our previous study^13^.

**Figure 1.**
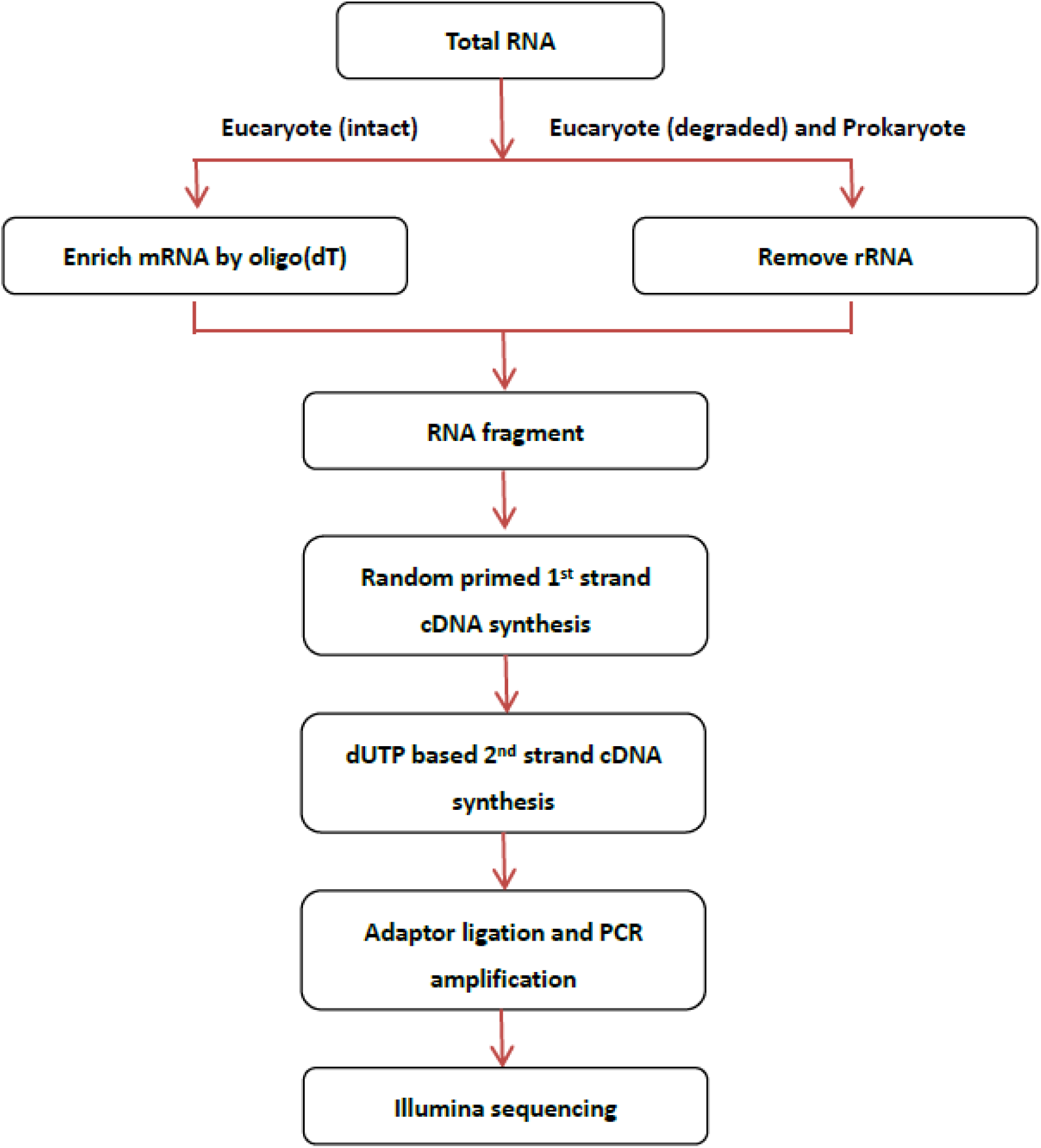
Flowchart of the RNA-sequencing Procedure.

### Statistical Analysis

Data are mean±SD for continuous variables. Categorical variables were presented inpercentages and numbers. Unpaired Student’s t-test and the chi-squared test were employed to compare continuous andcategorical variables, respectively. Fisher’s exact test was utilized for assessing GO enrichment and pathways, with p<0.05 considered of significance.R was employed for determining FPKM and performing other statistical tests such as hierarchical clustering of genes showing differential expression levels.

## Results

### Patient Characteristics

The major characteristics of the HF and non-HF groups showed no significant differences (**Table 1**). However, the HF group had elevated BNP amounts and LVEDD, and reduced LVEF, in comparison with the non-HF group.

**Table1.** Participants Characteristics.

|  | CAD with HF<br>(N=5) | CAD without HF<br>(N=5) | P value |
| --- | --- | --- | --- |
| Male, n(%) | 2(40) | 3(60) | 0.999 |
| Age, years | 67.8±5.0 | 60.8±8.1 | 0.139 |
| BMI, kg/m<sup>2</sup> | 23.8±4.6 | 25.0±5.5 | 0.718 |
| Diabetes, n(%) | 2(40) | 2(40) | 1.000 |
| Hypertension, n(%) | 4(80) | 5(100) | 1.000 |
| HbA1c, % | 6.85±0.87 | 7.00±1.81 | 0.871 |
| Total cholesterol, mmol/L | 3.67±3.07 | 3.60±1.52 | 0.965 |
| LDL-C, mmol/L | 2.34±2.52 | 2.24±1.46 | 0.941 |
| HDL-C, mmol/L | 0.84±0.53 | 0.70±0.12 | 0.580 |
| Triglyceride, mmol/L | 0.89±0.38 | 1.86±0.89 | 0.055 |
| Creatinine, µmol/L | 88.2±25.2 | 76.8±16.8 | 0.424 |
| Uric acid, µmol/L | 363.2±82.4 | 357.0±69.3 | 0.901 |
| BNP, pg/ml | 1795.4±1053.3 | 76.4±22.6 | 0.006 |
| LVEDD, mm | 60.0±5.9 | 47.5±5.2 | 0.007 |
| LVEF, % | 44.2±10.2 | 62.0±4.8 | 0.008 |

Notes: BMI, body mass index; HbA1c, glycosylated hemoglobin; LDL-C, low-density lipoprotein cholesterol; HDL-C, high-density lipoprotein cholesterol; BNP, brain natriuretic peptide; LVEDD, left ventricular end diastolic diameter; LVEF, left ventricular ejection fraction.

### RNA SequencingFindings

We performed RNA-seq analysis of ribosomal-depleted total RNAs extracted from EAT in CAD patientsof the HF and non-HF groups (n=5/group). In total, 2278 EAT circRNAs weredetected in the 10 CAD patients. The lengths of the EAT circRNAs detected were mainly within 2000nt (**Figure 2**). The numbers of exons per circRNA averaged 3 (ranging between 1 and 21). Single-exon circRNAs accounted for 9% (211/2278), while 2.7% (63/2278)of all circRNAs putatively had 10 exons or more. The circRNA hsa_circ_0087255 was the longest (21 exons). A total of 90% (2051/2278) circRNAs had average read counts of less than 10, while 0.5% (11/2278) had valuesexceeding 50. This indicated the latter group had highest levels in human EAT; these circRNAs corresponded to genes such as KIAA0182, RHOBTB3, HIPK3, UBXN7, PCMTD1, N4BP2L2, CFLAR, EPB41L2, FCHO2, FNDC3B and SPECC1 (**Table 2**).

**Table 2.**
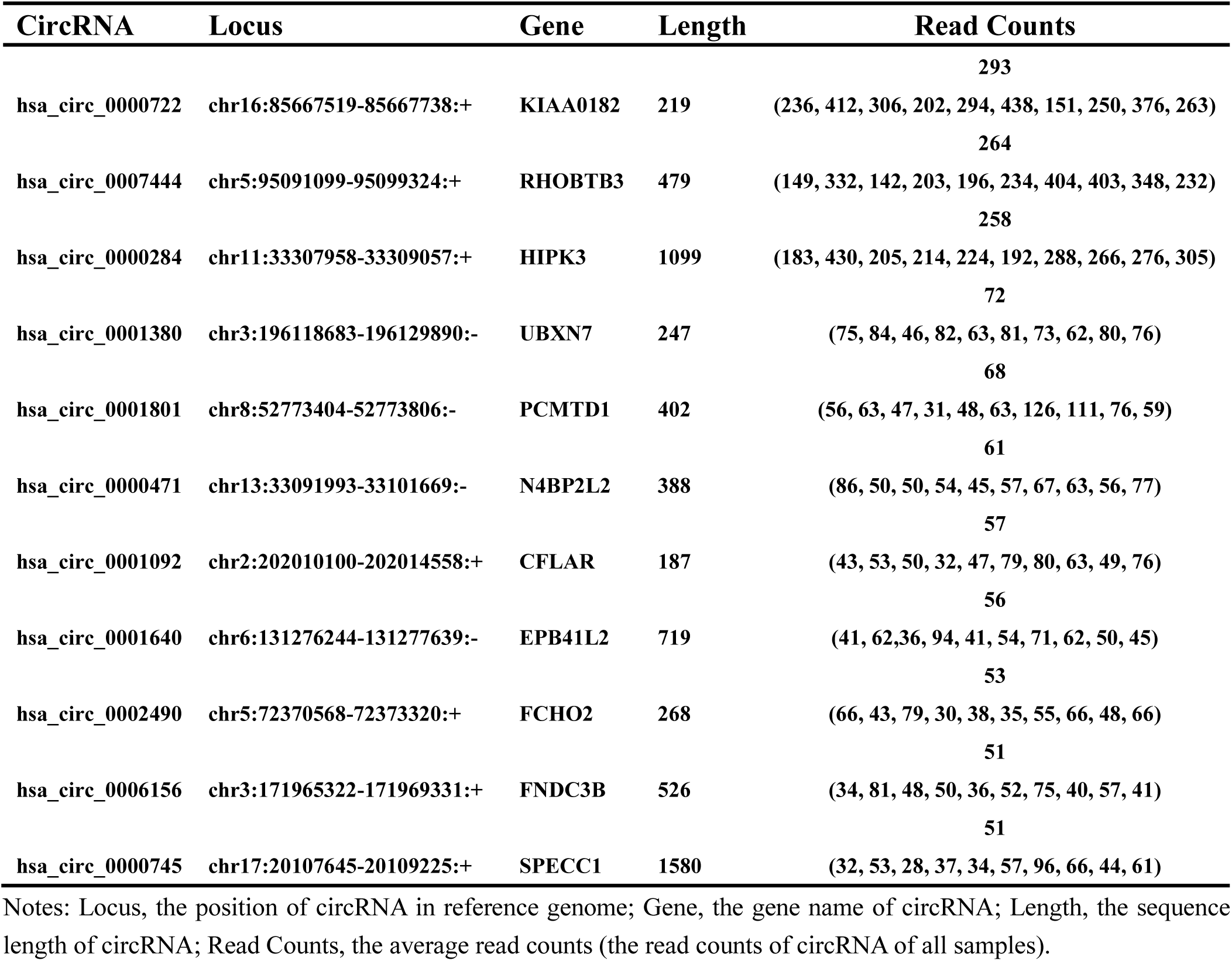
Top highest expressed circRNA in human epicardial adipose tissue.

**Figure 2.**
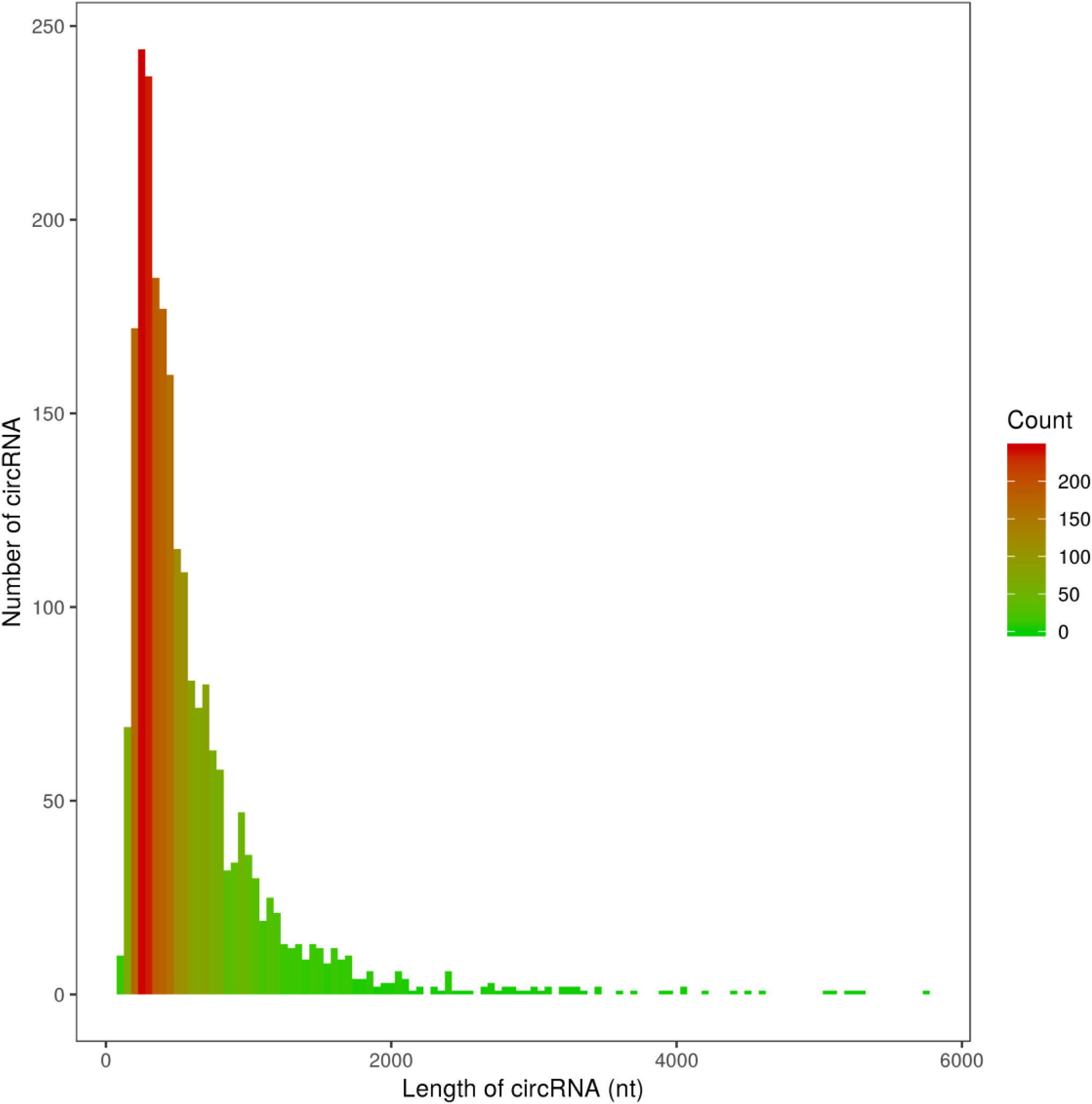
Length range of circRNAs in human epicardial adipose tissue.

Hierarchical clustering was carried out for grouping differently expressed circRNAs in CAD cases of the HF and non-HF groups based onexpression, and circRNA expression patterns in the HF group showed marked differences in comparison with the non-HF group. A total of 1240 circRNAs with significant level changes(p<0.05) were detected, with 561 upregulated and 679 downregulated (**Figure 3**).

**Figure 3.**
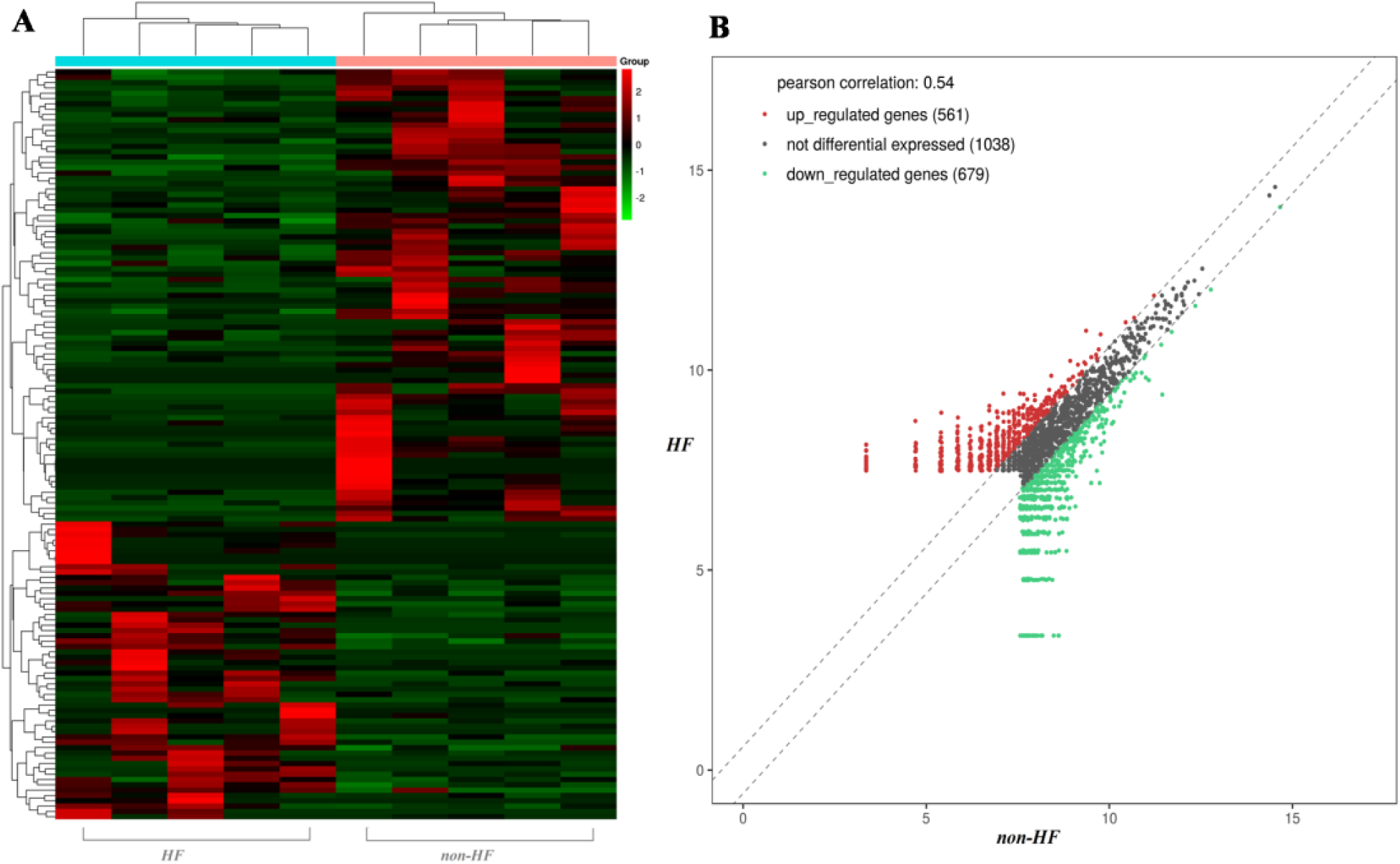
Hierarchical clustering (A) and scatter plot (B) of circRNAs with differential expression(p<0.05) in the heart failure (HF) and non-HF groups (red and green denoteupregulation and downregulation, respectively). A total of1240 circRNAs were found (561 upregulated and 679downregulated).

To explore the possible roles of circRNAs in EAT, we selected the circRNAs with substantially different amounts between the HF and non-HF groups (P<0.05;fold change >2)as potential novel biomarkersof HF; in total, they were 141, including 56 and 85 showing upregulation and downregulation, respectively (**Supplementary material, Table S1 and Table S2**). Among them, hsa_circ_0005565 had the highest fold change (27.4), and was highly expressed in all the 5 CAD patients with HF and lowly expressed in the non-HF group (**Figure 4**).

**Figure 4.**
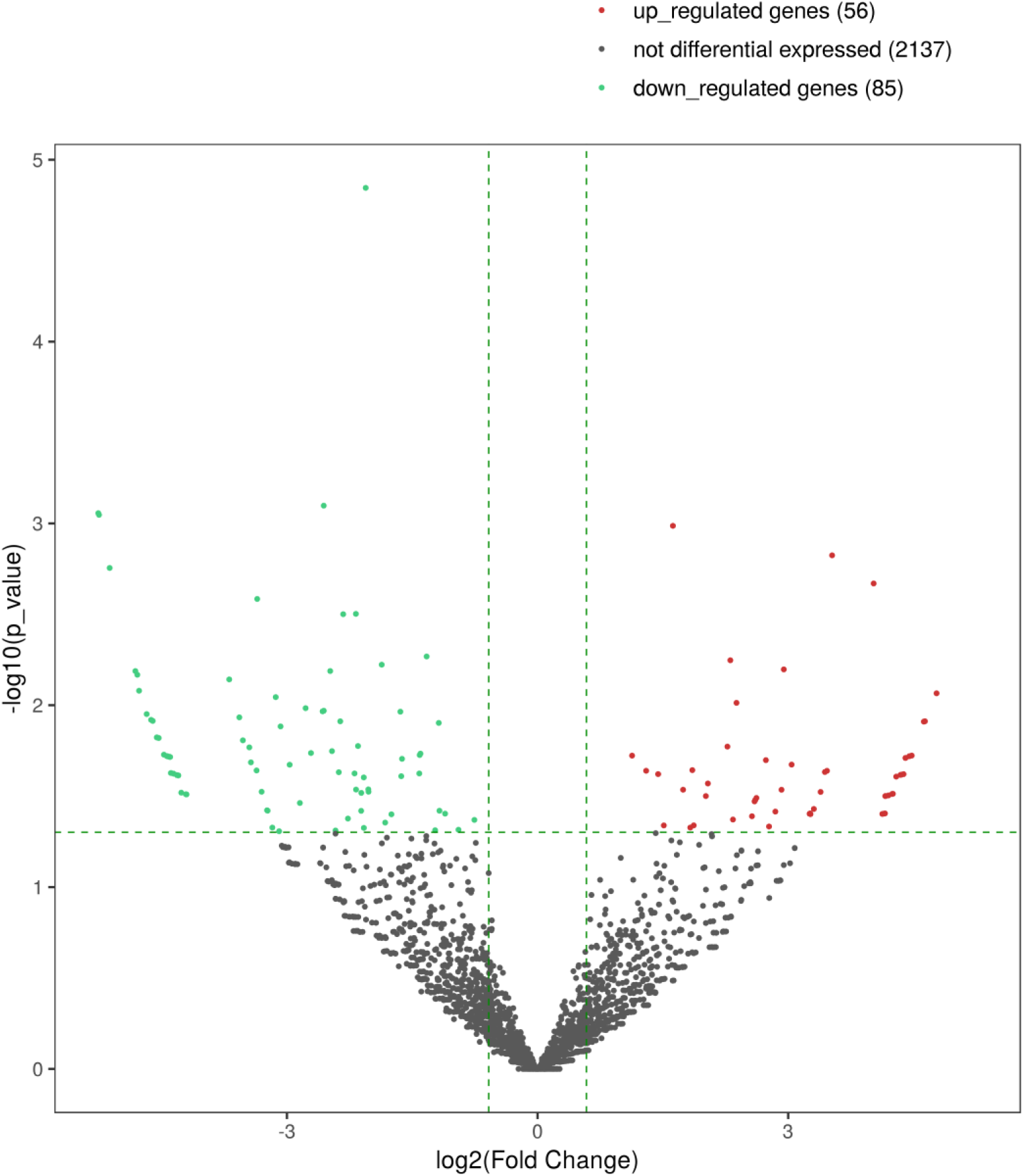
Volcano plot of circRNAs with differential expression(P<0.05;fold change >2) in the heart failure (HF) and non-HF groups (red and green denoteupregulation and downregulation, respectively). A total of 141 circRNAs were found (56 upregulated and 85 downregulated).

### Protein-coding genes associated with differentially expressed circRNAs

The majority of circRNAs have unknown functions. Therefore, GO and KEGG pathway analyses were carried out of circRNAs with differential expression between the HF and non-HF groups to determine the associated protein-coding genes. The 3 first enriched GO terms in order were positive regulation of metabolic process (GO: 0009893), positive regulation of cellular metabolic process (GO:0031325), and macromolecule metabolic process (GO:0043170). The first 3 KEGG pathways in order included insulin resistance (hsa04931), transcriptional misregulation in cancer (hsa05202), and platinum drug resistance (hsa01524).

## Discussion

Here, circRNA expression patterns in EAT of CAD cases were compared between the HFand non-HF groups. A total of 11 circRNAs were highly expressed in human EAT, including hsa_circ_0000722, hsa_circ_0007444, hsa_circ_0000284, hsa_circ_0001380, hsa_circ_0001801, hsa_circ_0000471, hsa_circ_0001092, hsa_circ_0001640, hsa_circ_0002490, hsa_circ_0006156 andhsa_circ_0000745).The circRNAs with differential expression in EAT among HF cases (including 56 upregulated and 85 downregulated circRNAs), as well as GO terms and KEGG pathways were also determined.

Expression profiling of circRNAs has been performed in various tissue types^14^. *In vivo*, circRNAs have been reported in cardiovascular tissues (right atrium, vena cava and heart) based on mouse or human GWAS data from blood cells^15, 16^. *In vitro*, circRNAs have been described in cardiomyocytes, cardiac fibroblasts, vascular smooth muscle cells, endothelial cells, macrophages, and blood monocytes^17-19^. A recent study^2^ assessed ribosome-free RNAs from human and mousecardiac specimens as well as human embryonic stem cell-derived cardiomyocytes by RNA-seq, detailedly revealing cardiac circRNA expression patterns, which provides a valuable reference for further circRNA analysis. However, circRNA expression profiles in human EAT remain largely unclear. Meanwhile, it is known that circRNAs are associated with different types of cardiovasculardiseases, such as HF^16^.

EAT is a link between metabolic disorders and HF. Its thickness, independently of BMI, is positively associated with left ventricularmass^20^. Epicardial fat amounts, without regard to metabolic status or CAD, correlates with impaired left ventricular function^21^and myocardial fibrosis^22^.ETA is considered to be involved in the pro-inflammatory polarization and fibrotic transformation occurring in HF^12, 22-24^. Similarly to other visceral adipose tissues, EAT secretes many molecules with exocrineand paracrine impacts on adjacent organs. These factors, released from EAT, might affect cardiac cell metabolism, as well as endothelial, arterial smooth muscle and inflammatory cell functions, causing HF.

In summary, the expression patterns of EAT circRNAs in CAD cases were identified, with special attention to those involved in HF. These findings also reveal potential HF biomarkers, which should be confirmed in further investigations.

## Author contribution statement

Meili Zheng and Lei Zhao carried out data analysis andmanuscript writing. Xinchun Yang performed study design.

